# Molecular dynamics study of differential effects of serotonin-2A-receptor (5-HT2AR) modulators

**DOI:** 10.1101/2025.03.27.645663

**Authors:** Jordy Peeters, Dimitri De Bundel, Kenno Vanommeslaeghe

**Author notes:** Corresponding author: (KV).

## Abstract

The serotonin-2A receptor (5-HT_2A_R) is an interesting target for drug design in the context of antidepressants that might have a rapid onset of action and/or be effective in treatment-resistant cases. The main challenge, however, is that the activation of this receptor can provoke hallucinations. Recent studies have shown that activating the receptor with certain (partial) agonists could potentially give rise to antidepressant effect without hallucinogenic side effects. Although substantial research has been done in this area, the atomistic details of this differential activation of the serotonin-2A receptor are not fully understood. In the present study we performed multiple atomistic molecular dynamics (MD) simulations on 5-HT_2A_R bound with an antipsychotic, two different potential non-hallucinogenic antidepressants and a hallucinogen to identify the receptor’s ligand dependent conformations. Overall, our findings suggest that modest 5-HT_2A_R activation would only yield antidepressant effects and hallucinations result from excessive activation. While modest activation through microdosing may be problematic on account of abuse potential as well as possibly narrow and patient-dependent therapeutic windows, modest activation through administration of a sufficiently weak partial agonist may offer a viable drug development pathway.

**Author Summary:** Activating the serotonin-2A receptor (5-HT2AR) with specific agonists is a promising strategy for the development of a new class of antidepressants. However, potent agonists such as LSD, DMT or psylocibin generally cause hallucinations. To aid the development of 5-HT2AR-targeted antidepressants that do not cause this side effect, we seek to gain a better understanding of the receptor by studying its activation mechanism. Specifically, we performed molecular dynamics simulations to explore how different drugs interact with 5-HT2AR, both in the absence and presence of an intracellular binding partner. This led to the identification of tentative intermediate states along the activation pathway, which could be linked to the ligands’ pharmacological properties. Our findings suggest that hallucinogens cause an excessive build-up of activated receptors, whereas carefully designed mild activators could lead to a new generation of antidepressants that do not induce hallucinations.

## 1 Introduction

The 5-HT_2A_-receptor belongs to the family of G protein-coupled receptors (GPCRs) and is a crucial component of the serotoninergic system, implicated in various mental health conditions including schizophrenia, bipolar disorders, ADHD, migraine, and depression-related disorders.[1–3] In treating schizophrenia and bipolar disorders, the 5-HT_2A_R is primarily targeted with atypical antipsychotics acting as either antagonist[4–10] or inverse agonist.[4,7,8,11–13] Conversely, the use of 5-HT_2A_ agonists is limited due to their propensity for inducing hallucinogenic effects. Since the ‘50s, depressive disorders are instead treated with TCAs, MAOIs, SSRIs and SNRIs,[14] which increase the concentration of endogenous serotonin in the synaptic cleft.

Clinical studies on hallucinogens remain limited, but investigations on 5-HT_2A_R’s agonists LSD, psilocybin and mescaline have shown rapid and long-lasting effects in the treatment of depressions, substance abuse and anxiety.[15–17] This suggests that direct pharmaceutical activation of the 5-HT_2A_-receptor could be helpful in the treatment of depression. Accordingly, several non-hallucinogenic 5-HT_2A_ agonists[18,19], including compounds IHCH-7086[20] and *(R)-*69,[21] have been reported to exert significant antidepressant but not hallucinogenic activity in mouse models.

The classic GPCR activation mechanism involves the formation of a multi-protein complex that includes the G-protein at the intracellular side of the activated receptor. In this environment, the G-protein is phosphorylated, setting in motion an intracellular signaling cascade. Eventually, the receptor is desensitized, which is thought to involve the blocking of the G-protein binding site by β-arrestin. However, β-arrestins have been shown to also exert influence on downstream signaling pathways, particularly those involving kinases such as the Src/AKT cascade.[22]

It is not fully understood why some 5-HT_2A_ agonists have antidepressant and/or hallucinogenic effects. Separate signals associated with G protein G_q_ and β-arrestin binding have been hypothesized to be involved. Specifically, Cao et al. synthesized a potential antidepressant non-hallucinogenic compound IHCH-7086.[20] This ligand shows high binding affinity towards 5-HT_2A_, no activation of the G_q_ pathway and showed a low maximum efficacy at the β-arrestin2 assay. However, the involvement of β-arrestin is still subject to debate, as demonstrated by Kaplan et al.’s partial agonist (*R*)-69, which is biased to G protein pathway activation (*E*_max_=87% relative to 5-HT) without any β-arrestin2 recruitment, yet is also reported to have non-hallucinogenic antidepressant effects in mouse models.[21] In another study, Wallach et al. demonstrated that an agonist’s hallucinogenic potential correlates with its efficacy for the Gq pathway. Specifically, they found that ligands with an E_max_ below 70% (relative to 5-HT) did not induce the head-twitch response in mice, a known predictor of hallucinogenic activity in humans.[23] Taking these studies together seems to suggest that partial agonists with low efficacies can give rise to antidepressant effects without psychedelic properties regardless of their activation pathway. The present work seeks to contribute to the understanding of these differential effects through a detailed MD study of the receptor itself.

### 1.1 Molecular activation mechanism

The general activation mechanism of GPCRs has been studied extensively. On a macroscopic scale, an outward movement of Transmembrane Helix 6 (TM6) occurs upon activation, accompanied with an inward movement of TM3 and TM7 as shown in Fig. 1A. Note that the Ballesteros-Weinstein nomenclature is employed in further sections.[24] At the level of individual interactions between side chains, the activation of GPCRs involves breaking the ionic lock of the E/DRY motif between residues Glu^6×30^ of TM6 and Arg^3×50^ of TM3. Additionally, the hydrophobic interaction between Ile^3×46^ and Leu^6×37^ in these helices also breaks. This allows TM7 to rearrange itself so its NPxxY motif (Asn^7×50^, Pro^7×51^ and Tyr^7×53^) interacts with a hydrophobic residue of TM3 (Ile^3×46^). In turn, the side chain of the tryptophan “toggle switch” (Trp^6×48^) rotates and TM6 bends at the level of the Phe^6×44^ residue, forming an interaction between the Ile^3×40^ and Phe^6×44^ residues of the P^5×50^I^3×40^F^6×44^ motif. Furthermore, agonists can create a hydrogen bond with Ser^5×46^ resulting in an inward bulge in TM5.[25] Interestingly, this serine is substituted by an alanine residue in both rodent 5-HT_2A_ receptors and human 5-HT_2B/C_ receptors.[26] Therefore, this residue is thought to be involved in differences in ligand affinity through different species, sub-type selectivity binding, binding kinetics[27] and biased signaling.[28] It should be mentioned that these microswitches can be toggled independently, i.e. that ligands can alter the energetics for each switch differently.[29]

**Figure 1:**
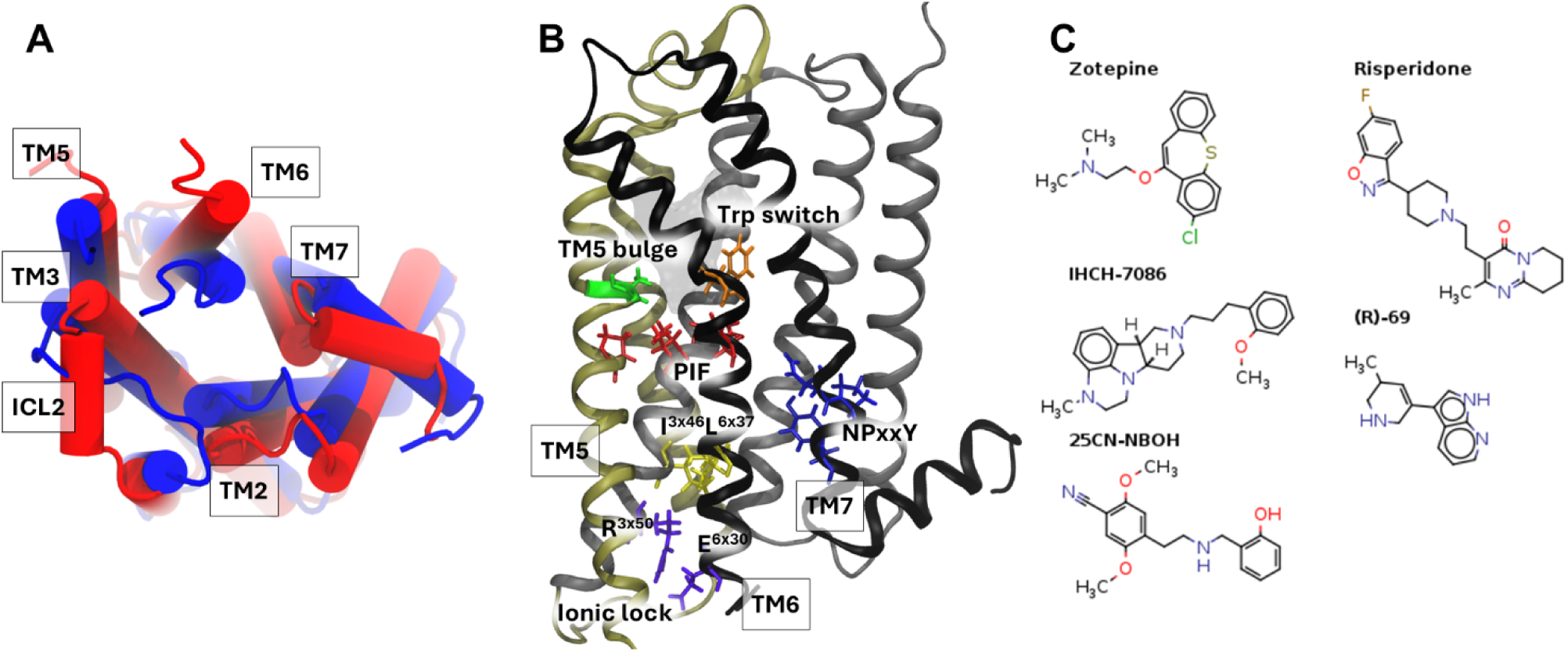
(A) The intracellular side of 5-HT_2A_R. In red the conformation in its active form and in blue its inactive one. (B) The serotonin-2A receptor with important side chains highlighted. (C) An overview of the ligands used for MD simulations.

Throughout the GPCR family, the highly conserved tyrosine Y^5×58^ and Y^7×53^ (the latter is part of the NPxxY motif) forms a mediated hydrogen network which is believed to stabilize the active state.[30] More specifically for 5-HT_2A/B/C_ receptors, residue Phe^6×41^ further stabilizes these side chain conformations through π-π stacking whereas this residue in other GPCRs contains a hydrophobic non-aromatic side-chain.[28]

### 1.2 Modulation of 5-HT_2A_

Previous studies[25,31–34] have shown that multiple combinations of “on” and “off” microswitches are likely relevant in the activation pathway and/or the differential response to different types of GPCR ligands. To survey the resulting landscape of states in the specific case of the serotonin-2AR, we performed MD simulations with 5 different ligands; zotepine, an antipsychotic antagonist, risperidone, an antipsychotic inverse agonist, IHCH-7086, a β-arrestin biased partial agonist, (*R*)– 69, a G-protein biased partial agonist (both with retained hallucinogenic properties) and lastly, 25CN-NBOH, a β-arrestin biased partial agonist with hallucinogenic properties. These structures are displayed in Fig 1C. As it became clear during the study that an intracellular binding partner is necessary for the receptor to reach its active state, we performed each MD simulation in the absence and presence of an intracellular G protein construct. This allowed us to correlate the various pharmacological profiles of the selected ligands with the dynamics of the 5-HT_2A_R by mapping their different effects on key microswitches as well as on large-scale helix movements during MD simulations.

## 2 Materials and methods

Five distinct experimental structures of the 5-HT_2A_R, each complexed with ligands exhibiting different pharmacological effects, were used to explore the conformational landscape by means of atomistic molecular dynamics (MD) simulations. Eleven simulations were performed using these templates. Table 1 provides an overview of these simulations along with the pharmacological properties of the ligands.

**Table 1:** Overview of performed MD-simulations. [a] The templates of simulations 6 – 11 contain a G-protein construct. [b] The hallucinogenic ligand was replaced in the PDB with the corresponding ligand.

| Simulation | Ligand<sup>[a]</sup> | pharmacological effect | Drug class | PDB entry template | Simulation time (μs) |
| --- | --- | --- | --- | --- | --- |
| 1 | Zotepine | Antagonist | Antipsychotic | 6A94 | 2.8 |
| 2a | None | Basal |  | 6A94 | 4.8 |
| 2b | None | Basal |  | 6WHA | 4.1 |
| 3 | Risperidone | Inverse agonist | Antipsychotic | 6A93 | 3.2 |
| 4 | IHCH-7086 | β-arrestin biased partial agonist | Potential non-hallucinogenic antidepressant | 7WC9 | 4.2 |
| 5 | (R)-69 | G-protein biased partial agonist | Potential non-hallucinogenic antidepressant | 7RAN | a) 3.4<br>b) 3.6 |
| 6 | 25CN-NBOH | β-arrestin biased partial agonist | Hallucinogen | 6WHA | 3.0 |
| 7 | Zotepine + G | Antagonist | Antipsychotic | 6WHA<sup>[b]</sup> | a) 1.9<br>b) 1.8 |
| 8 | None + G | Basal |  | 6WHA | a) 2.4<br>b) 2.1 |
| 9 | IHCH-7086 + G | β-arrestin biased partial agonist | Potential non-hallucinogenic antidepressant | 6WHA<sup>[b]</sup> | a) 2.1<br>b) 1.3 |
| 10 | (R)-69 + G | G-protein biased partial agonist | Potential non-hallucinogenic antidepressant | 7RAN | a) 3.0<br>b) 3.2 |
| 11 | 25CN-NBOH + G | β-arrestin biased partial agonist | Hallucinogen | 6WHA | a) 1.8<br>b) 1.2 |

### 2.1 System preparation

The X-ray and cryo-EM structures listed in Table 1 were downloaded from the RCSB PDB databank (http://www.rcsb.org/pdb).[35] and prepared for MD using CHARMM-GUI.[36] PDB IDs 6A93 and 6A94 were a crystallographic dimer, of which only chain A was retained. In PDB IDs 6A93, 6A94 and 7WC9 the apocytochrome b562RIL was fused to the third intracellular loop (ICL3) for crystallographic reasons. Manually removing this fusion protein resulted in a chain break between residues 265 (in TM5, the 5^th^ transmembrane helix) and 313 (TM6). PDB ID 7WC9 contained mutations S372N, S162K and M164W and its residues 181 to 187 were unresolved.

These artefacts were corrected using the CHARMM-GUI PDB manipulator.[37–39] In the case of 6WHA, the rudimentary loop modeling functionality did not succeed for 6WHA’s missing residues 69-83 of TM1 and residues 348-353 of extracellular loop 3 (ECL3); the coordinates of those amino acids were copied from 7WC9 after alignment of the surrounding secondary structure elements. The cryo-EM structures 6WHA and 7RAN contained a mini G_q_ protein[40] complexed with the single-chain variable fragment (scFv16); these were left out during the system preparation of simulation 5 and 6. The mol2 files of zotepine, risperidone, IHCH-7086, (R)-69 and 25CN-NBOH were obtained by building the ligands in Avogadro and were subsequently protonated at their most basic N-atom using OpenBabel[41]. This resulted in protonation of the basic amine, which is important because the experimental structures suggest a polar hydrogen bond with Asp 155 of the receptor. The topology and parameter files of the ligands were generated automatically with CGenFF based on the mol2 during the PDB manipulation step.[42] In addition, the PDB file of 7WC9 included monoolein molecules. Accordingly, a topology file for monoolein was constructed manually from CHARMM lipid force field[43] atom types; see “available data”. The N-and C-termini of the protein (including the ICL3 chain break, where present) were capped as acetylamide and methylamide groups, resp.

The “preprocessed” proteins resulting from the above procedure were solvated in a rectangular periodic membrane system using CHARMM-GUI.[42–48] By means of trial and error, the box dimensions and the thickness of the water layer were chosen so the distance between the receptor and its images was at least 24 Å. The orientation of the receptor in the membrane was based on the Orientations of Proteins in Membranes database.[49] Ultimately, the X and Y dimensions of the box were both set to 75 Å and the water thickness to 13.5 Å. Next, a heterogeneous bilipid membrane layer was selected which represents the biological environment of the receptor. Specifically, based on the lipid composition of synaptic vesicles,[50] the membrane was built with 57 POPC (1-palmitoyl-2-oleoyl-sn-glycero-3-phosphocholine), 18 POPS (1-palmitoyl-2-oleoyl-sn-glycero-3-phospho-L-serine), 8 PSM (1-palmitoyl-2-stearoyl-sn-glycero-3-phosphoethanolamine), 12 CER160 (ceramide 160), 52 POPE (1-palmitoyl-2-oleoyl-sn-glycero-3-phosphoethanolamine), 4 POPI (1-palmitoyl-2-oleoyl-sn-glycero-3-phosphoinositol) and 42 cholesterol molecules. The replacement method was used for building the membrane. After a ring penetration check, the membrane was solvated in a 0.15 M KCl solution using the Distance Monte Carlo algorithm implemented in CHARMM-GUI.

For simulations 7-11, a small construct representing the parts of the G protein that influence the receptor was assembled by manipulating the PDB files of 6WHA and 7RAN. Specifically, residues 25-39, 69-86 and 222-246 of the mini-G_q_ protein in chain B were retained (at their original positions). All N-and C-termini *that resulted from breaking up the chains* were capped with an acetylamide and methylamide group, respectively; note that this does not include the actual C terminus of the chain as used in the Cryo-EM structure (resid 246). The retained residues are displayed in Fig S1. In order to conserve the geometry of the original mini-G_q_, we defined distance restraints on a selection of atoms given in Table S1 along with their force constants. After positioning the resulting construct next to the receptor based on structure alignment, the proteins were solvated in rectangular periodic membrane systems as described above (but resulting in a slightly larger box).

### 2.2 Equilibration steps and molecular dynamics

The system was heated and simulated with NAMD 2.14[51] using the standard protocol and simulation parameters from CHARMM-GUI.[43,52] Briefly, a 10000-step minimization was followed by three 250 ps equilibration steps and three more 500 ps equilibration steps, each at 303.15 K but with gradually weakening restraints. In the subsequent production phase, a time step of 2 fs was used in conjunction with restraints on the hydrogen atoms using the SHAKE algorithm. The pressure of the system was kept at 1 atm using NAMD’s Nose-Hoover Langevin-piston barostat with a piston period of 50 fs and a piston decay time of 25 fs. A constant temperature was maintained using Langevin dynamics with a damping coefficient of 1 ps^-1^. Non-bonded interactions were smoothly switched off over 10 to 12 Å using a switching function for the electrostatics and a force switching function for the van der Waals interactions (as recommended for using the CHARMM force field in NAMD). The long-range electrostatic interactions were calculated using the particle-mesh Ewald method with a mesh size of 1 Å and a 6^th^ order spline interpolation.

### 2.3 Trajectory analysis

Interpretations, measurements and analysis was performed using VMD 1.9.3[53] and MDAnalysis 2.4.2[54] including the helanal package[55] to analyze helix bends and twists. Principal component analysis (PCA) was performed on the time series of the Degrees of Freedom (DOF) described in Table 2 (see Results and Discussion) using the Scikit-learn Python package.[56]

**Table 2:** PCA variables and their explanations. For more detailed information on their definition, we refer to Table S2 in the SI.

| PCA attribute label | Internal degree of freedom |
| --- | --- |
| TM26 | Outward movement TM6 |
| TM37 | Inward movement TM3 and TM7 |
| Ionic lock | Ionic lock breaks between R3x50 and E6x30 |
| TM36 extra | Extracellular distance TM3 and TM6 |
| TM36 intra | Intracellular distance TM3 and TM6 |
| TM46 | Movement of TM6 relative to TM4 |
| TM47 | Movement of TM7 relative to TM4 |
| TM56 extra | Extracellular distance between TM5 and TM6 |
| TM56 intra | Intracellular distance between TM5 and TM6 |
| TM67 | Intracellular distance between TM6 and TM7 |
| I3x46 L6x37 | Hydrophobic interaction Ile169 and Leu325 |
| I3x40 F6x44 | distance Ile163 and Phe332 of the PIF motif |
| F34x56 R3x50 | Distance Phe186 (ICL2) and Arg173 (ionic lock) |
| P34x50 I3x40 | Distance Pro180 (ICL2) and Ile163 (PIF) |
| TyrTyr | Distance Tyr254<sup>5x58</sup> and Tyr380<sup>7x53</sup> |
| S5x46 G7x41 | Distance S242 to G369 to measure TM5 bulge |
| TM5 twist | Average helix twist near TM5 bulge |
| TM6 bend | Average helix bend of TM6 |
| $\chi_1$ W6x48 | $\chi_1$ dihedral angle of Trp336 (Trp switch) |
| $\chi_2$ W6x48 | $\chi_2$ dihedral angle of Trp336 (Trp switch) |
| $\chi_1$ F6x44 | $\chi_1$ dihedral angle of Phe332 (PIF motif) |
| $\chi_2$ F6x44 | $\chi_2$ dihedral angle of Phe332 (PIF motif) |
| RMSD inact | RMSD relative to inactive conformation |
| RMSD NPxxY inact | RMSD NPxxY relative to inactive conformation |
| RMSD PIF inact | RMSD PIF relative to inactive conformation |
| RMSD active | RMSD relative to active conformation |
| RMSD NPxxY activ | RMSD NPxxY relative to active conformation |
| RMSD PIF active | RMSD PIF relative to inactive conformation |
| RMSD ICL2 activ | RMSD of ICL2 relative to active ICL2 helix |

The 2D probability density plots in §3.4 were constructed using SciPy’s kernel density estimator.[57] More specifically, the Boltzmann inverted populations obtained from the different MD simulations are shown in Figs 5 and 6. While these should not be interpreted as converged free energy profiles, the logarithm makes the large range of population densities more manageable as well as expressing them in more intuitive units of energy.

## 3 Results and discussion

### 3.1 Principal component analysis

Given the plethora of microscopic modulations of 5-HT_2A_R in response to different ligands, we attempt to systematize the changes in the aforementioned macro– and microswitches. Known relevant DOF in the receptor are listed in Table 2, along with their definitions and measurement methods. To find patterns in the high-dimensional space of these DOF, it was subjected to PCA, retaining the first four principal components (PCs) for closer interpretation.[56] Simply put, the first PC captures the most variance, with subsequent PCs explaining the remaining variance that was not already captured by earlier PCs. in this fashion, three PCAs were performed on three conformational data sets: (1) all the trajectories in this study (Fig S2), (2) only the trajectories in the absence of the G protein (Fig S3) and (3) only the trajectories in the presence of the G protein (Fig S4).

When considering the PCA loading plots derived from the union of all trajectories (Fig 2A), the hydrophobic interaction (I^3×46^ L^6×37^), ionic lock and the attributes related to the movement of TM6 are all situated along the positive axis of in PC1, indicating that they constitute the major source of correlated motions. In addition, the TM3, TM7 and TM5 helix movements appear along the negative PC1 axis, indicating an anticorrelation. This is in excellent agreement with the consensus activation mechanism wherein they play important roles and indirectly suggests that relevant active and inactive states were sampled. For the sake of readability, we displayed only a selection of DOF in Figs 2A-C; full loading plots for PC1 through PC4 are presented in Figs S2-4. Perhaps more surprisingly, DOFs related with the tryptophan switch (the dihedral angle χ_1_ of Phe332^6×44^, the χ_2_ angle of Trp336^6×48^ and TM5 twist) are described by the second principal component, indicating that their dynamics are largely independent of PC1. The distance between TM3-7 (anti)correlate in both PCs. The importance of each attribute in PCs 3 and 4 are given in Figs S2-4. Though the variance between the attributes in these subsequent PCs become less correlated, one could carefully interpret PC3 as the component which is related with the helix movements of 5-HT_2A_Rs extracellular side. No clear interpretation of PC4 was found.

**Figure 2:**
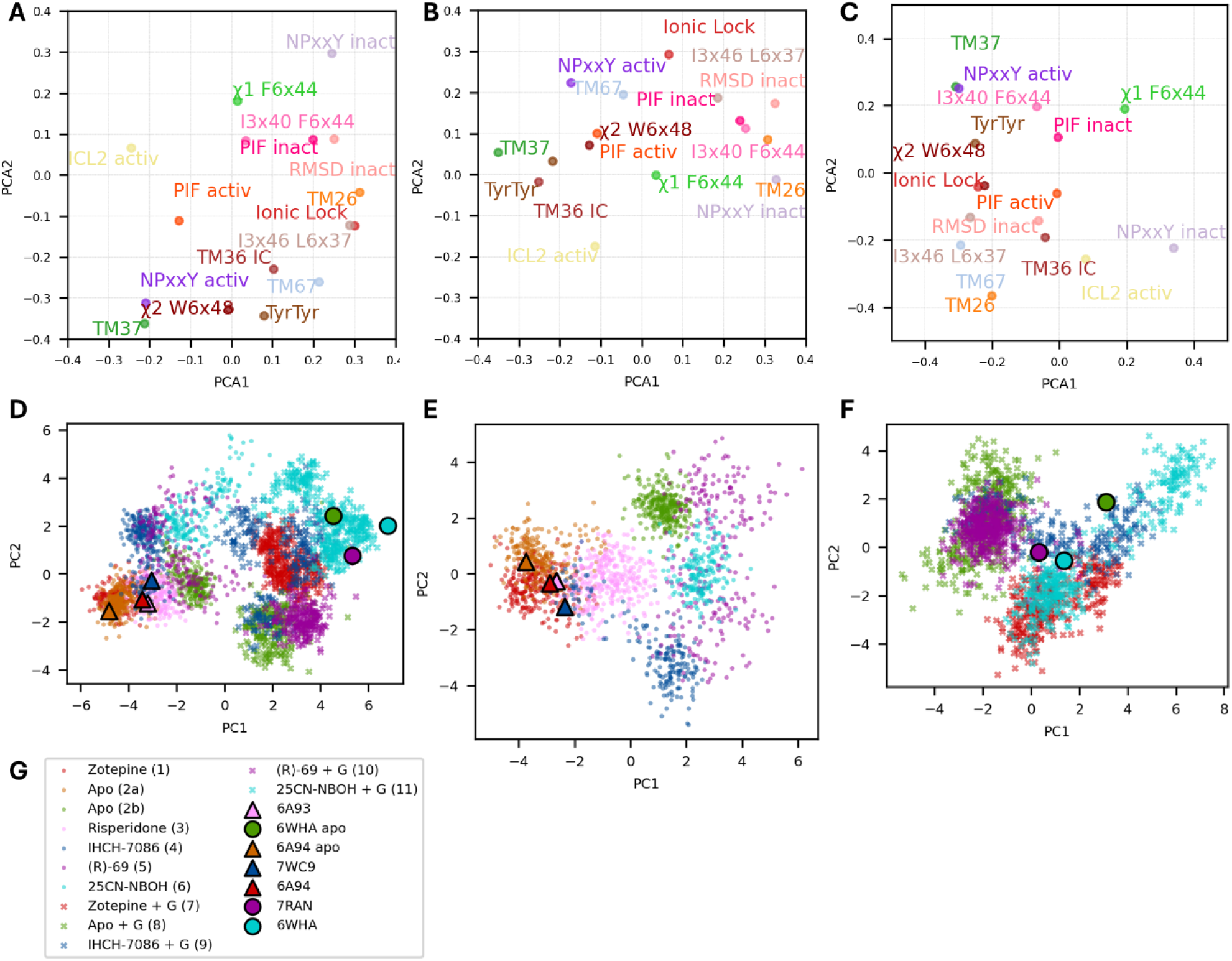
PCA results for (left) simulation with and without G protein (1-11), (middle) without (1-6) and (right) with G protein construct (7-11). Top row: partial loading plots results; some DOF are omitted for clarity but all are present in Figs S2-4. Bottom row: PC1 and PC2 for all trajectory frames that were included in the respective PCA.

From our MD simulations we know that the G protein is necessary for the receptor to reach its fully active conformation (discussed later) meaning that the position of TM6 is more inward in the resting state. Accordingly, the outward movement of TM6 is a hallmark event even upon activation and the importance of the associated DOFs stands out in our PCA. However, when the PCA attributes of the receptor in the absence (Fig 2B) and in the presence (Fig 2C) of an intracellular binding partner are compared, the relative importance of this hallmark event drops because the ligand alone is not able to push TM6’s outward shift. This indicates that ligand-dependent factors modulate these other DOFs such that the TM6’s outward shift becomes more energetically feasible. In Fig 1B and Fig 1C, and in more detail Figs S3-4, attributes related to TM7 and its NPxxY motif dominate PC1. This suggests that the differential nature of the ligands has a direct impact on these degrees of freedom, even if a full transition between an active and inactive state (with binding/unbinding of the G protein) is not observed over the course of the simulations.

The trajectories of the MD simulations listed in Table 1 are represented by the two PCs in Fig 2D-F. Figs S2-4 contain analogous plots for PC3 and 4. The circles and triangles in the plots of Fig 2D-F represent the receptor’s X-ray or CryoEM state, respectively, whereas the clusters of dots and crosses represent frames of different trajectories. The first principal component can be seen as representing the activated nature of the receptor conformation with respect to intracellular signaling, starting from the left hand side with inactive receptor states (circles and dots) lacking an intracellular binding partner progressing rightward to states with the G protein (triangles and crosses). Conversely, the second component distinguishes between partial active states and may also correlate with the activating nature of the ligand present in the binding site. Looking in more detail, the effect of the ligand on the respective MD ensemble is also apparent, starting on the left hand side with the antagonist zotepine, followed by (R)-69 (purple) and IHCH-7086 (blue), which are believed to be non-hallucinogenic agonists. Further rightward are the clusters found where the receptor is bound with the hallucinogenic compound 25CN-NBOH (cyan). Lastly, the apo simulation 2a, starting from the inactive template 6A94, forms a distinct cluster from apo simulation 2b, which started from the active template 6WHA. Specifically, simulation 2a (orange dots) seems to be biased towards its initial inactive state (orange triangle). Conversely, simulation 2b started from an active state (green circle); upon removal of the G-protein and ligand, it quickly falls back to a *partially* active state (green dots) similar to the (R)-69 bound complex (purple dots), but remains quite distinct from the fully inactive state. This suggests the existence of a kinetic barrier that makes the conversion between these two states a rare event on our MD time scales.

### 3.2 Removal of the G protein

Time scales in MD simulations are generally too short to reliably observe the transition of 5-HT_2A_R to an active state. However, studies have shown that an intracellular binding partner is necessary for at least some GPCRs to maintain their active conformation.[30,58] By removing the G protein, we can study the *deactivation* mechanism instead. In this context, Dror et al.[32] observed three distinct states during the MD simulation of βAR in which an intracellular nanobody stabilizing the receptor’s active state was removed. Specifically, a plot of the RMSD of the NPxxY motif of an inactive conformation versus the TM3-TM6 distance showed three distinct clusters of conformations. The first cluster is positioned around the crystal structure. A second cluster is formed after the rearrangement of the NPxxY motif near the end of TM7. From this state, a last cluster is obtained when the distance between TM3 and 6 decreases. In an attempt to replicate these findings for the 5-HT_2A_R, similar plots were constructed in Fig S5 for simulations 2b, 5 and 6, in which the mini-G_q_ protein was removed from their corresponding PDB templates. In Fig S5, the black star on the 2D represents the state captured in the cryo-EM structure. In the first row are the TM3-6 and the RMSD of the NPxxY values plotted, revealing two clusters: one with a high RMSD value and with low RMSD values. Overall, these plots suggest that the NPxxY motif initially shifts toward a more inactive conformation, followed by a decrease in the TM3-6 distance before it subsequently increases again. An exception occurs in simulations 5a and b with ligand (R)-69 after the removal of the G protein. This simulation was run twice and transitioned to an inactive or partially active conformation along two distinct pathways. This would seem to imply that the rearrangements of the helices do not necessarily have to happen in a specific order.

The same analysis was also performed for two other degrees of freedom: the intracellular TM2-6 and TM3-7 distances as defined in Table S1. This pair of distances was chosen as representative for the shape of the intracellular binding surface. Indeed, as shown in Fig 1A, helices 2, 3, 6 and 7 form a square delineating this binding surface. Combining the information from the different experimental structures, a global anticorrelated motion must occur along the perpendicular TM6-2 and TM3-7 axes during the transition from an inactive apo state to a fully active state containing the mini-Gq protein and an activating ligand. However, our MD simulations indicate that two motions tend to happen sequentially rather than in a concerted fashion, as can be seen in Fig S5 H-J.

### 3.3 Microswitches, structural motifs and side chain conformations

The outward movement of TM6 is often associated with a bend in the helix near residues W6×48 and F6×44. Figures 3A and 3E show the distribution of TM6 bend angles. The top row represents simulations without the G protein construct (1-6; Fig 3A-C). In the simulations with the G protein construct (7-11; Fig 3E-G), the distributions are shifted to the right, indicating that the G protein induces a greater bend in TM6. Moreover, the histograms with G-protein have longer “tails” towards larger bending values, especially for the apo simulation (green curve in Fig 3E). This indicates that the presence of the G protein construct generally facilitates the bending motion of TM6.

**Figure 3:**
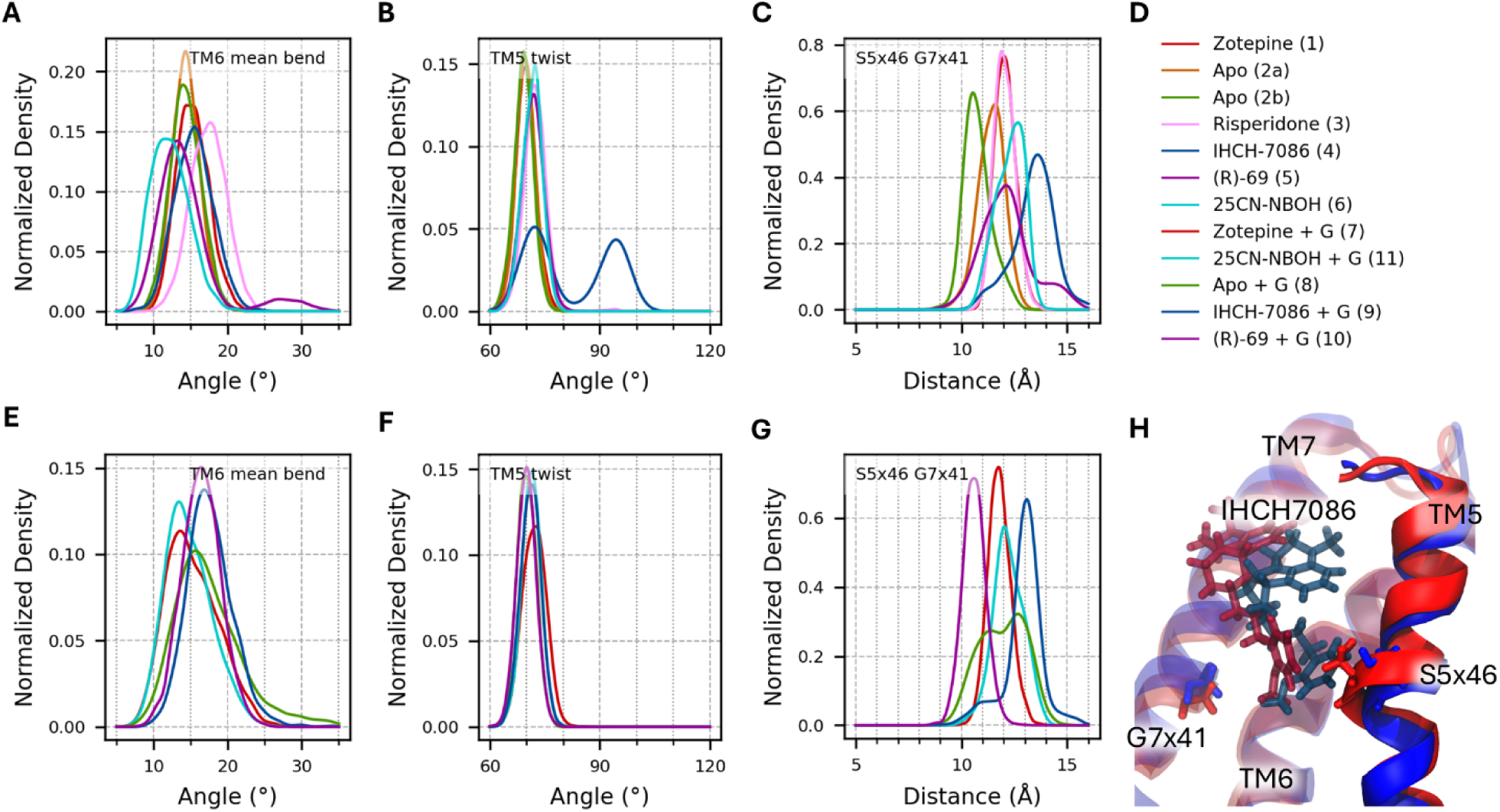
Normalized histograms are shown for simulations 1-6 without the G protein (top row) and simulations 7-11 with the G protein construct (bottom row). A and E: Average TM6 bend between residues 329 – 339. B and F: TM5 twist angles. C and G: Distance between closest heavy atom of S242^5×46^ and G369^7×41^. D: Legend of the histogram plots. H: Visualization before (red) and after (blue) the bulge in TM5 vanished in simulation 4 (IHCH7086).

A closer examination of the simulation distributions without the G protein (1-6; Fig 3A) reveals that TM6 has a greater degree of bending in the simulation with risperidone (pink) compared to the other simulations. The least bending occurs in the simulation with the hallucinogen (cyan), despite this ligand being considered the most activating. However, the TM6 bend itself is not a primary contributor to the variance in the PCA. Instead, the main explanator of PC2 are the residues Trp336^6×48^ and Phe332^6×44^ which are located near the bend. Fig 4 displays the χ_1_– and χ_2_-angles of these residues. In simulations with (R)-69 (5b) and 25CN-NBOH (6), a transition of the χ_2_-angle of the tryptophan switch between 110° (“off” state; Fig 4F) and 50 ° (“on” state; Fig 4G) could be observed. Moreover, the presence of the G protein populates the “on” states in the apo simulation (8) and the 25CN-NBOH bound ones (11).

**Figure 4:**
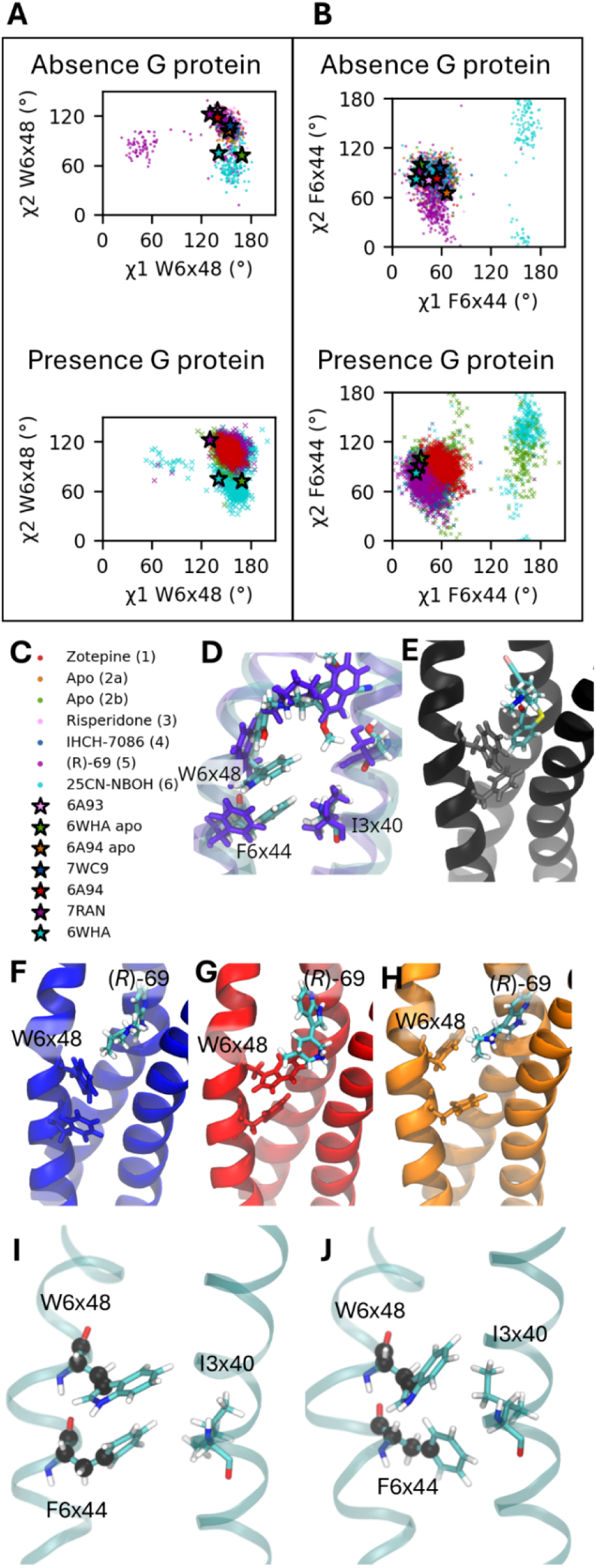
χ1 and χ2 angles of (A) the Trp336^6×48^ switch and (B) the Phe332^6×44^. (C) Legend of A and B. (D) Overlap between an “on” tryptophan switch (cyan) with the hallucinogen and an “off” one (blue) with (R)-69. (E) Blocked “Trp-switch” with antagonist zotepine. (F) Trp-switch in its “off” state and with (R)-69. (G) Trp-switch in its “on” state. (H) Trp in an alternate conformation with (R)-69. (I-J) Phe332^6×44^ in a gauche and anti-like conformation.

As suggested by Wallach et al.[23] and Liao et al.[33], different rotamers of Trp336^6×48^ may play a role in biased signal transduction. Their studies indicate that when the switch undergoes a further downward movement in the binding pocket—corresponding to smaller χ2 values—it promotes β-arrestin transduction. Consistent with their study, an unusual transition of the tryptophan switch occurs in simulation 5b (and briefly in simulation 10), with ligand (R)-69, and in simulation 11b with 25CN-NBOH. In these cases, instead of moving further down the binding pocket, the indole moiety of W^6×48^ rotates away so that χ_1_ assumes values between 30 to 60° (plotted in Fig 4A, and side chains illustrated in Fig 4H). This specific indole conformation is only observed in simulations with the G protein biased ligand (R)-69 and in the receptor-G protein complex in combination with a potent agonist like 25CN-NBOH. Therefore, this specific state depicted in Fig 4H could be an interesting target for future docking studies aimed at identifying additional G protein-biased ligands with low efficacies.

Figure 4D displays an overlap between the psychedelic ligand 25CN-NBOH (cyan) and the G protein biased candidate antidepressant (R)-69 (blue). Although they occupy roughly the same binding pocket, 25CN-NBOH possesses an additional aromatic ring which seems to be incompatible with the inactive state of the tryptophan switch, thus forcing it into its “on” state, whereas (R)-69 biases the tryptophan residue to an alternative “on” position from where its inactive state remains accessible. This observation might help explain the pharmacological differences between the two agonists. Therefore, agonists that extend less into the bottom of the ligand-binding pocket might promote a G protein-biased rotamer of Trp336^6×48^. Conversely, the antagonist zotepine penetrates much deeper into the binding pocket, blocking this switch and preventing interactions between the side chains in the connector region (Fig. 4E).

Additionally, Fig 4B displays χ_1_-χ_2_-plots for Phe332^6×44^, which is part of the highly conserved PIF motif. The χ_1_-value indicate that the benzyl moiety can adopt a gauche-like (Fig 4I) and antiperiplanar-like (Fig 4J) conformations. Notably, the antiperiplanar conformation is observed only in the presence of a strong activating ligand (e.g., the hallucinogen in simulations 6 and 11) or intracellular binding of mini-Gq (as in simulation 8). While in this anti conformation (Fig 4J), it becomes clear from that the phenyl group is allowed to rotate freely around its axis (see the χ_2_ values in Fig 4B and their time series in Fig S6). Taken together, these findings suggest that the tryptophan switch disrupts the π-π stacking between its aromatic group and Phe332^6×44^. This disruption may promote the anti conformation of the Phe332^6×44^ side chain, allowing the phenyl ring to rotate freely. This effect might initiate a cascade of conformational changes throughout the receptor’s intracellular side, biasing it toward binding an intracellular partner.

As briefly mentioned in the introduction, while residue S242^5×46^ does not interact directly with a ligand in 5-HT_2A_ receptors, it might have a role in biasing the signaling pathway of GPCRs. Fig 3B and 3F shows the (average) twist angle near this residue. In addition, the closest heavy atom distance between S242^5×46^ and G369^7×41^ is another way to measure the inward bulge of this residue which is displayed in Fig 3C and 3G.[59,60] While this bulge most often arises by an agonistic H-bond donor in other GPCRs such as β_2_ adrenergic and dopamine receptors, it is interestingly enough always present in our simulations. One exception occurs in simulation 4 when the β-arrestin biased ligand, IHCH-7086, is bound. After 1500 ns, the bulge in TM5 vanishes as shown in Fig 4H (blue) (time series are shown in Fig S7). When looking at the distance distribution between residues S242^5×46^ and G369^7×41^, different distributions for the three different agonists (purple, blue and cyan) are observed, especially when bound to the intracellular partner. We speculate that the bulge itself does not necessarily play a role in biased agonism, but that the distance between S242^5×46^ and G369^7×41^ might be a distinguishing factor.

### 3.4 Large scale movements

Since the outward motion of TM6 does not happen simultaneously with the inward motions of TM3 and TM7 during activation, different partially-active conformations can be proposed. To further investigate the populations of these different states, 2D probability density plots of TM6-2 vs TM3-7 are displayed in Fig 5.

**Figure 5:**
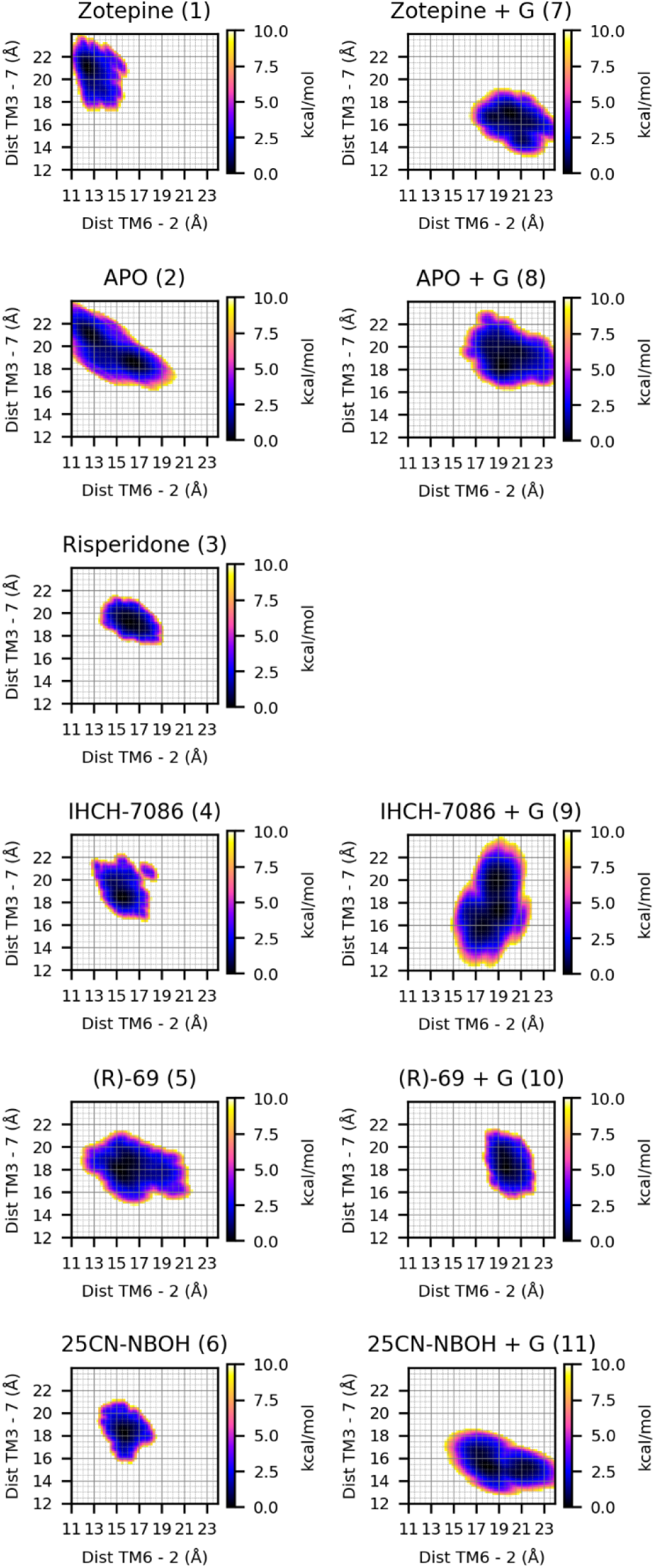
Boltzmann-inverted density plots for TM6–2 and TM3–7 axes, presented in free-energy units (kcal/mol). The left side shows simulations conducted without the G protein construct, while the right side displays simulations with the G protein construct present.

When comparing the non– and G-bound systems, a notable shift in the density to higher TM26 distances is observed. This shift indicates that the G protein is essential for TM6 to achieve its full outward position, consistent with the PCA results and the discussion in §3.2. In addition, when looking at the plots for the hallucinogen bound receptor (6 and 11), it can be observed that the minimum is shifted slightly downward, meaning that TM3 and TM7 are shifted inward in response to the presence of the G protein. Combing these observations, it would indeed appear that the hallucinogen-G protein-bound complex is located the farthest along an activation pathway consisting of an increase in TM2-6 and decrease in TM3-7 distance. Conversely, if we compare the location of this minimum with the other simulations, we can carefully assume that the other G protein-bound complexes are in a partially activated state, since either TM6 is shifted more inward, TM3 and TM7 shifted outward or both. The only exception to this trend is the system containing both the antagonist zotepine and the G protein. Indeed, if we compare the minima between simulation 7 and 11, we observe that the location is roughly the same, indicating that after docking zotepine into the G-protein-bound active conformation, the receptor fails to assume a less-active state on the time scale of the simulation, in contrast with the G-protein-bound simulations with a partial agonist or no small-molecule ligand at all. While this may easily be a shortcoming of the computational time constraints, an alternative explanation would be that zotepine hinders the conformational freedom of the receptor. More specifically, it seems plausible that zotepine would increase the free energy of one or more intermediary conformations in the activation mechanism, thus inhibiting both deactivation and activation. Indeed, in the absence of the G protein there is a trend away from the upper-left corner of the graph with increasingly activated ligands, while the simulation with zotepine displays a compact distribution corresponding to a fully deactivated state. We further speculate that zotepine accomplishes this by sterically hindering the unraveling of a network of interactions that holds TM6 in place in both the activated and the deactivated conformation. This is supported by Fig 4E, where zotepine hinders the tryptophan switch and the interaction between F^6×44^ and I^3×40^ of the PIF motif.

In simulation (8), upon removal of the hallucinogen 25CN-NBOH, the population density shifts toward higher intracellular TM3-7 distances, again indicating that both the G protein and the hallucinogen are necessary for the receptor to remain in the conformational basin corresponding to the active state. When the G protein-biased or β-arrestin biased partial agonist is bound to the receptor (simulations 9 and 10), it is observed that they have a different effect on the TM7-3 distance. Conversely, their non-G protein bound counterparts (simulations 4 and 5) seem to position TM6, TM3 and TM7 in a more “active”-like position when compared with zotepine, especially for the G protein biased agonist. In addition, we found that upon G protein activation, the hallucinogen 25CN-NBOH populates a structural different state than the other agonists. Comparing its conformational landscape (simulation 11) with the partial agonists (9 and 10) leads us to speculate that the ideal β-arrestin biased receptor conformation would feature a TM37 distance close to 15 Å and a TM26 distance around 18 Å (see red in Fig. 6). In this hypothesis, TM26 should be somewhat higher (around 21 Å) for G protein signaling, which would be potentiated by a further decrease in TM37. Combined with 25CN-NBOH’s high absolute binding affinity for the receptor, this would make it plausible that it causes hallucinations by inducing excessive populations of activated 5-HT_2A_ while at the same time being β-arrestin biased. It would also explain why both IHCH-7086 and (R)-69 are weak agonists while being β-arrestin and G protein biased, respectively.

Another 2D density plot is given in Fig 6 in which the TM26 and TM37 distances of all simulations is included.

**Figure 6:**
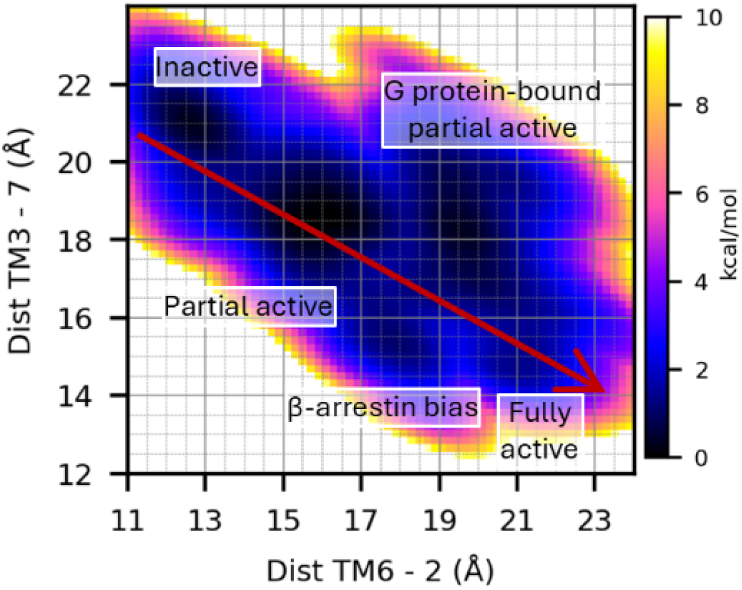
An overall density plot including the TM6-2 and TM3-7 distances of simulation 1-11. It shows the distinguishable basins (labeled by the 5-HT_2A_’s state) modulated by different ligands. A putative “axis of activation” is drawn in red along the TM6-2 and TM3-7 axes.

In summary, after some equilibration, the simulations without G protein largely assumed an inactive state, especially along the TM6-2 axis – however, they showed a clear tendency *in the direction of* the active state for increasingly agonistic ligands, which is most pronounced along the TM3-7 axis as can be observed from the distinguishable basins in Fig 6. Similarly, the simulations with G protein largely remained in an active state (again, especially along the TM6-2 axis) but showed a clear tendency *in the direction of* the inactive state for increasingly inactive or antagonistic ligands, at least along the TM7-3 axis. The only exception to the latter observation is the simulation with zotepine, which biased the conformational ensemble toward an inactive state without G protein, but toward an active conformational ensemble with G protein, despite it being known as an antagonist. This may simply be a consequence of insufficient sampling, but if not, the simulation results suggest that binding of zotepine to the inactive state might hinder the (microscopic) conformational changes necessary to “unlock” the large-scale motion of the transmembrane helices. Regardless, the global trend is consistent with the hypothesis that an activating ligand by itself does not make 5-HT_2A_ transition to its fully activated conformation, but rather adds a bias to the conformational energetics such that the binding of the G protein *concurrent* with the transition to an active state (presumably near the lower right of Fig. 6) becomes energetically more favorable. Indeed, this mode of action has been proposed on the basis of computational and experimental data for the β2-adrenergic receptor[25,32] and might be more widespread in the GPCR family. Going one step further, our results regarding the G protein versus β-arrestin biased ligands are in excellent agreement with the observation by Weis and Kobilka that despite their differential pharmacology “these [different] agonists likely share allosteric transmission mechanisms”.[25] Taken together, this leads us to hypothesize that hallucinations result from excessive populations of activated 5-HT_2A_ and that a non-hallucinogenic antidepressant effect could be achieved by *a sufficiently mild activating ligand*. This may imply an energetic effect that *weakly favors* the “fully activated” conformation (i.e. a *quantitative* effect), either by binding relatively weakly to the receptor (like the endogenous ligand 5-HT) or by binding strongly but having a modest activating influence of the conformational energetics. Alternatively, a mild activating ligand could induce a strong bias in the receptor’s conformational energetics, but toward a “non-ideal” conformation that only weakly enhances the intracellular binding of the G-protein (which we would call a *qualitative* effect). Such a conformation might lie between the inactive and fully active conformation or correspond to the β-arrestin biased conformation (which traditionally would lead to desensitization). Either way, the above distinction between a “qualitative” and “quantitative” effect is difficult to make when studying 5-HT_2A_ in isolation because the conformational difference between the tentative G protein biased and β-arrestin biased conformations is modest, so that Hammond’s postulate predicts that both effects will often occur together. This equally implies that this distinction may be of lesser concern in practical efforts to design non-hallucinogenic 5-HT2A antidepressants.

## Conclusions

The results of this study suggest that intracellular binding of the G protein is necessary for the 5-HT_2A_ receptor to assume a fully activated conformation exhibiting the common outward movement of TM6 in GPCRs. Indeed, we propose that the role of activating ligands is not to *directly* induce this outward movement, but to alter the conformational energetics of the receptor such that the (possibly but not necessarily simultaneous) adoption of an activated conformation and intracellular binding of the G protein becomes more favorable, in line with the β2-adrenergic receptor. This variant of the GPCR activation mechanism would naturally give rise to a range of partial agonists that differentially modulate the equilibrium between 5-HT_2A_R’s resting and G protein-bound active conformation. We further hypothesize that, *while strong activators are known to induce hallucinations*[23]*, a sufficiently weak (partial) agonist – either by virtue of weak binding or weak activation upon binding – might be able to avoid this undesirable side effect while still functioning as an effective antidepressant*. It is worth noting that this hypothesis leaves the question open whether such a weak or partial agonist should be G-protein or β-arrestin biased.

Indeed, both pathways likely have a shared transmission mechanism at the level of the receptor, as proposed by Weis and Kobilka[25] and supported by the present results. We argue in §3.4 that both intracellular effects will typically occur together because of Hammond’s postulate. Therefore, the question becomes more academic, and is tied to ongoing debates regarding the intracellular signals that give rise to antidepressant effects and which types of 5-HT_2A_ targeted therapies would be effective. It should be noted in the latter context that, if a small population of artificially activated receptors indeed yields an antidepressant effect, then precisely tailored small doses of a full agonist could theoretically be a viable alternative to a suitable partial agonist. While this “microdosing” route has been the subject of considerable interest and debate,[61–63] even if effective, it raises serious pharmacological and regulatory concerns. Overall, our results support the general idea of directing investigative efforts toward (weak) partial agonists, such as (*R*)-69 and IHCH-7086.

## Acknowledgements

This work was supported by the Vrije Universiteit Brussel (VUB) through research council (OZR) starting funds (OZR2893, to Kenno Vanommeslaeghe). The supercomputing resources and services used in this work were provided by the VUB and the VSC (Flemish Supercomputer Center, project 2023_100), the latter being funded by the FWO and the Flemish Government.

## Supporting information

**S1 Table**: Definition of the collective variables (CVs) in order to preserve the geometry of the G protein construct.

**S1 Figure**: Intracellular side of 5-HT_2A_ with the G protein construct.

**S2 Table**: Attributes used for the PCA.

**S2-4 Figures:** PCA loadings plots.

**S5 Figure**: 2D time series plots for simulations 2b, 5 and 6.

**S6-7 Figures**: Evolution of the specific dihedral angles.

